# ROCA: An Ontology to Describe and analyse Tool Use and Tool Making

**DOI:** 10.1101/2022.09.29.510031

**Authors:** Pierre R. Mercuriali, Carlos Hernandez Corbato, Geeske H.J. Langejans

## Abstract

To contribute to current discussions on the complexity of extinct and modern non-human and human primate tasks, we explore if ontologies are helpful as a means of representing tool tasks. We thus hope to help illuminate how primates approach tool tasks internally, such as breaking open a nutshell between a wooden anvil and a stone hammer to access its nutritious contents or fishing for termites with a leaf midrib. Ontologies are widely used in domains such as linguistics, medicine, archaeology, and cultural heritage, to help experts organize, reason on, and discover new knowledge in their field. We build a novel ontology, ROCA (from ‘roca’ - - rock, in Spanish) with which we can describe instances of tool use in a formal and uniform manner, including well-known primate ethograms and chaînes opératoires. We will see that representing tool use and tool making with an ontology provides a **uniform, unified, acentric, dynamic, and human-readable way to handle knowledge obtained from literature and to perform knowledge discovery**. We build a representative corpus of 75 articles and books on primate and hominin tool use and tool making. We then extract and give semantic structure such as taxonomical relationships to relevant vocabulary, both manually and automatically, using NLP text mining techniques. We then show how the ontology can be used to discover new knowledge related to tool use and tool making.

## 1. Introduction

Tool behaviours in our hominin family are generally considered an expression of advanced cognition and inter and intra species comparisons are key in debates on cognitive requirements for tool behaviour. Tool use and tool making in extinct and modern non-human and human primates are commonly described as a sequence of actions. However, currently it is difficult to compare and analyse these tool use and tool making sequences. Here, we explore the potential of ontologies to create a unifying method and framework to study tool behaviours.

Archaeological, anthropological, and primatological literature contains data from observations and reconstructed behaviours (experimental archaeology). These can be organised in *chaînes opératoires* (Carvalho et al. 2008, Bar-Yosef and Van Peer 2009) and textual narratives. These descriptions are the foundation of the comparative study of ancient and modern people, other primates, and other species. However, there is no common scheme of description within and across different fields. The little common ground in the categories used sometimes makes the representation schemes incompatible. It is, for example, impossible to compare the techno-steps in Perreault et al. 2013 and Kozowyk et al. 2017, as their categories and assumptions do not match. Thus, when behaviours are compared it may remain superficial. For example, comparisons of technological procedures may rely on measurements, such as the number of steps in a behavioural sequence, the length of a process or the procedural units. Deep analysis through computer-aided reasoning and statistics is generally impossible because there is only one (simple) matching concept. In addition, due to the variety of representations, aggregation of behavioural data across existing literature and own observations is complex, again also hampering complex analysis of large datasets.

### Let us consider the case of the *chaînes opératoires*

A *chaîne opératoire* is a descriptive and analytical tool to represent activities. The assumption is that every technical activity follows a certain chronological development and involves transformations that are either visible or invisible (Techniques & Culture 2019). Robert Cresswell gives the following formal definition: “[a *chaîne opératoire*] corresponds to a sequence of operations that transform raw material into a product, whether the latter is an item to be consumed or a finished tool” (Cresswell 1976). Furthermore, a *chaîne opératoire* always has a direction towards a known outcome (Martinelli 1985). However, concepts, connections, and assumptions, are generally not made explicit. The latter makes it difficult to compare *chaînes opératoires* made by different researchers who may use different terms and emphasise different components and outcomes: in its most general form, *a chaîne opératoire* needs not follow specific syntactic guidelines. This also makes it difficult for computers to manipulate the data and reason automatically over it. In addition, although there is an enormous increase in digital data on primate tool use with the democratisation of video recordings and ethology software, there is no unified structure from which the research community can draw to upscale their datasets. *Chaînes opératoires* have been criticized for the above reasons and as such they also cannot be used as a unified analytical framework (Bar-Yosef and Van Peer 2009). What if they, and other useful representation schemes, could all be represented within the same computer-friendly data structure?

### Ontologies offer a potential solution

An *ontology* is, in its most basic representation, a taxonomy, that is, a system of classification based on ranking and hierarchical relations. For example, in Figure 1, we represented a taxonomy of a subset of primates relevant to tool use and tool making. The different concepts are related with the sole relation ‘isA,’ also referred to as *conceptual subsumption*, which expresses for example that *JapaneseMacaque* is a *Macaque*. However, in addition to a taxonomy, concepts in an ontology can be linked within a directed labelled graph. Labelled edges in the graph express a multitude of relations, and concepts can have multiple links. For example, in Figure 2, we represented an instance of a chimpanzee inserting a stick into a tree. In the representation, the relation *actor* has for domain the concept *InsertingStick* and for codomain the concept *Chimpanzee*, to indicate that in this situation we describe a chimpanzee inserting a stick. Compare with Figure 1: the only relation that is available in a taxonomy is the conceptual subsumption ‘isA’, which indicates that the *InsertingStick* is an *Inserting action*.

**Figure 1:**
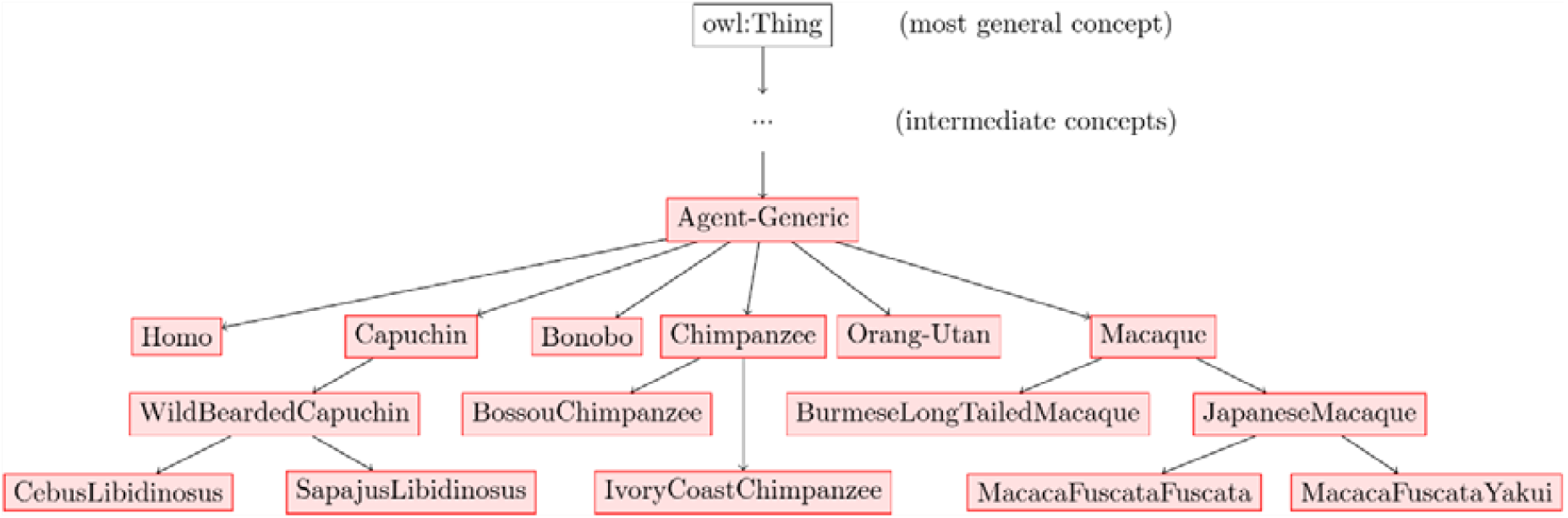
graph representation of part of the phylogenetic relations in the ROCA ontology. An arrow from concept A to concept B indicates that ‘B is an A,’ as in ‘a *JapaneseMacaque* is a *Macaque*.’

**Figure 2:**
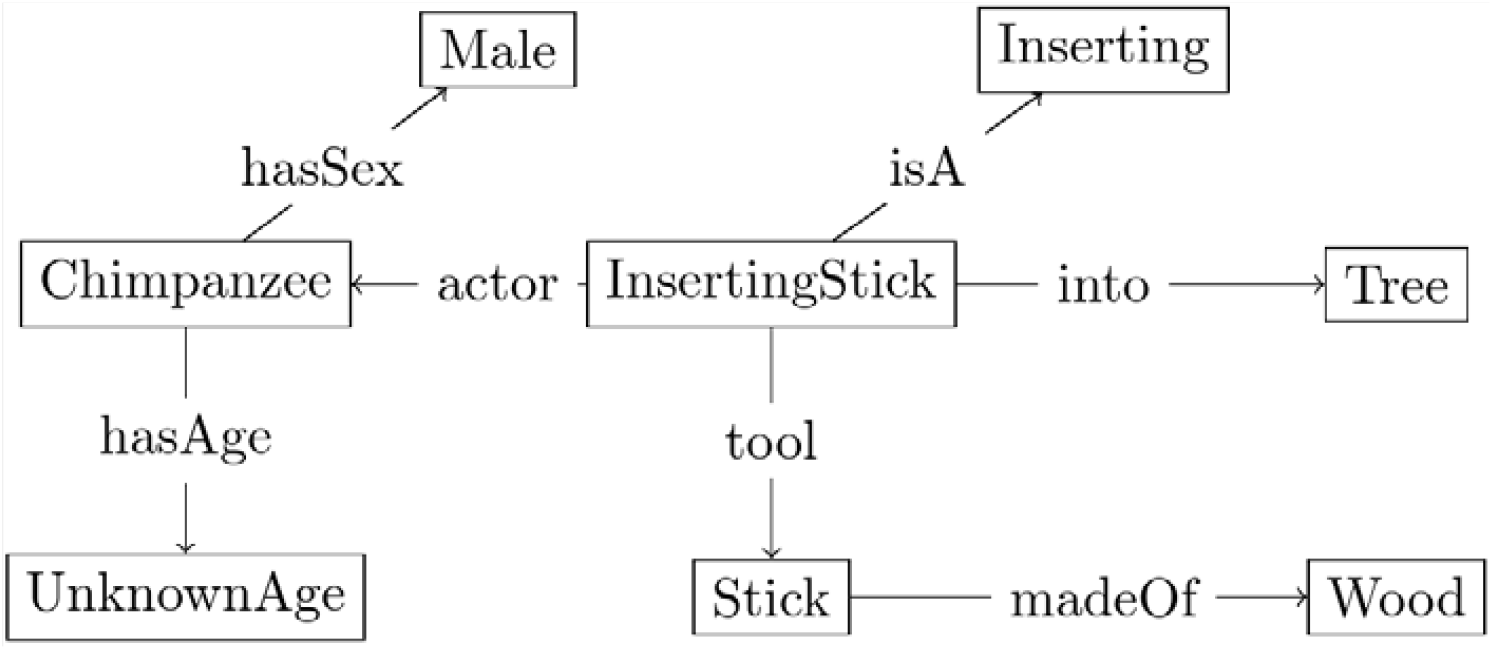
Concept graph representation of a chimpanzee inserting a stick into a tree.

Large datasets from the literature allow us to obtain taxonomic structure and new representations of knowledge with non-directed labels (Gruber 1995). Going even further, actual concrete descriptions and reconstructions of behaviour can be managed directly within the ontology using predefined concepts and relations.

Ontologies and ontology languages such as Web Ontology Language (OWL) are already widely used to facilitate knowledge discovery and reasoning. For example, the ORDO ontology deals with rare diseases, and it provides a structured vocabulary, capturing relationships for example between diseases and genes. ORDO provides a unified language for health professionals and a framework for the computational analysis of rare diseases (Orphanet, INSERM 1997). Other (bio)medical ontologies capture genetic (Osborne et al. 2009) and anatomical data (Mungall et al. 2012). Linguistics ontologies can take the form of multilingual semantic networks, like BabelNet (Ehrmann et al. 2014) and WordNet (Miller et al. 1990). Ontologies are also used in archaeology and cultural heritage studies: for example, the CIDOC Conceptual Reference Model (Doerr 2003) and the ARIADNEPlus infrastructure (Niccolucci and Richards 2013). The first provides a shared semantic framework to map any cultural heritage information published in any heritage discipline such as the museum world, commercial archaeology, and archives. The ontology unites otherwise disparate semantic or geographic data into one resource that can also be queried in further studies.

**Here we set out to first create a unifying ontology for primate tool use and subsequently we tested its usefulness to handle data obtained from the literature. We show that with an ontology we can overcome some of the research issues we have outlined above**. Moreover, we demonstrate that it can used to obtain new knowledge with computer-aided reasoning.

## 2. Materials and Methods

We set up to build an ontology for non-human primate tool use and tool making. We describe our building process, and the textual sources that give us the important concepts related to tool use and tool making.

Ontologies were first defined by Gruber in (Gruber 1993, page 1) as “an explicit specification of a conceptualization” (also see Olivares-Alarcos et al. 2019). In case of ROCA, we conceptualized *individuals* that are the things the ontology is about, *classes* that are properties the individuals may satisfy, *functions* to identify and relate individuals, and *axioms* to state what is true in the ontology (Staab and Studer 2009). Ontologies can be classified by how general the nature of the knowledge they aim to conceptualize is. A so-called *upper ontology* like the Basic Formal Ontology (BFO, Arp, Smith, and Spear 2015) and the Descriptive Ontology for Linguistic and Cognitive Engineering (DOLCE, Borgo 2009) focus on general concepts such as *Event, State, Object, Thing*. They include general relations, such as *parthood* or *participation*, and are not restricted to a specific discipline. On the other hand, a *reference ontology* has more specific concepts and relations and is relevant to a specific discipline. ROCA, because it aims to capture tool use and tool making among primates, thus aims to be a *reference ontology* for tool behaviours.

We built the ontology from published descriptions and reconstitutions of tool use behaviour y primates and early hominins (for information on maintenance and technical details of ontologies see Arp, Smith, and Spear 2015; Fernández-López, Gomez-Perez, and Juristo 1997). We started off with part of the pre-existing reference ontology SOMA (SOcio-physical Model of Activities) by (Beßler et al. 2020), based itself on the upper ontology DOLCE. SOMA is a useful base because it already contains physical concepts related to tool use, such as grasping a food or tool items or describing agents and their capabilities.

An agent is understood as an individual that acts (from Latin *agens, acting, doing*) and perceives its environment – for more information on the notion of agents, see (Russel and Norvig 2020, chapter 2). SOMA also contains a taxonomy of these concepts and thus already provides a starting structure to our ontology. First, we extracted relevant vocabulary from raw text of a corpus of primate literature. Second, we added semantic structure to the vocabulary, taking inspiration from the structure already given in SOMA, and thereby creating our ontology. Third, we used the structured vocabulary to add instances of tool use from raw text to the ontology (for more detail on knowledge-discovery processes see Roth et al. 1983; Gruber 1995). In Figure 3, we give an overview of the ontology-building process.

**Figure 3:**
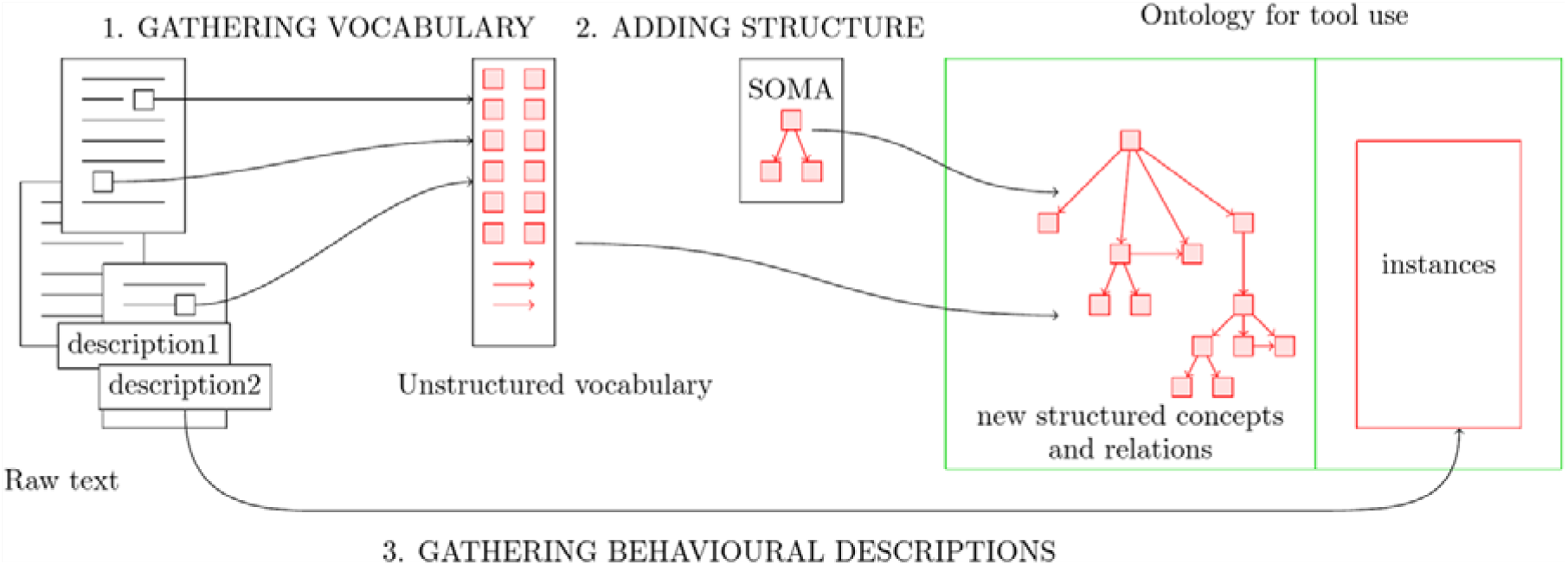
Overview of ontology-building process. The steps are aided by automatized scripts.

### 2.1. Data collection: gathering relevant vocabulary

The first step in building the ontology is to gather the important concepts and relations between them that we want to capture. For example, “cracking,” “stone,” “hammer,” “anvil,” “Coula nut,” “chimpanzee” is important vocabulary. We describe the process of building an unstructured vocabulary list from raw text, summarized in Figure 4.

**Figure 4:**
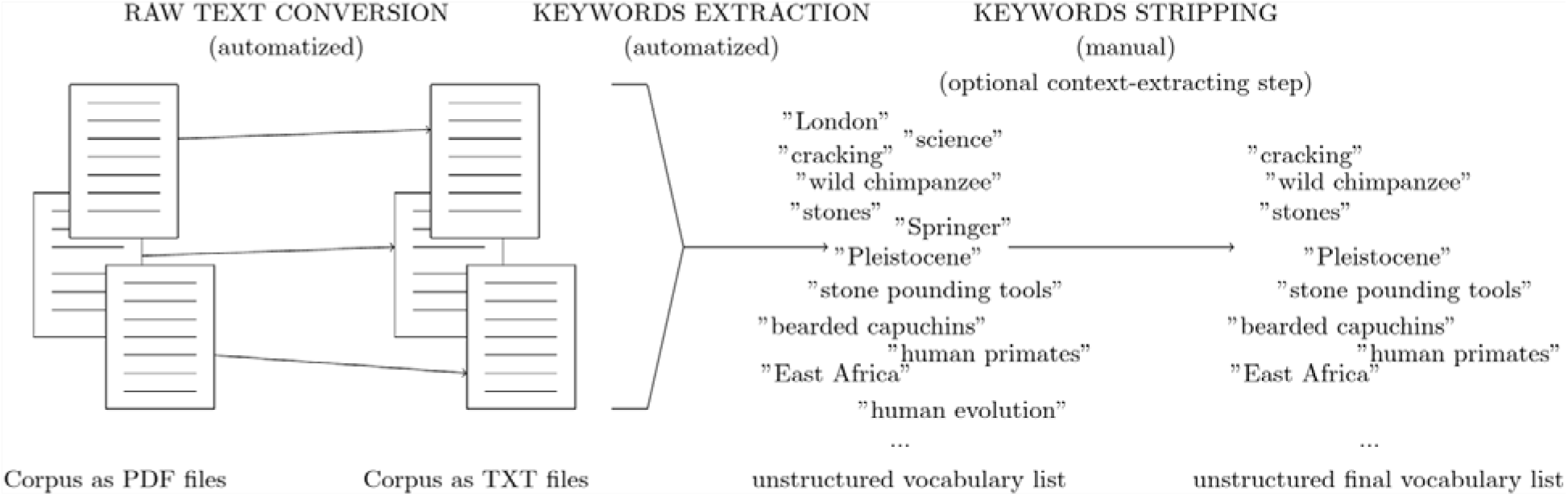
Overview of the extraction of keywords from a set of articles in the commonly used PDF format. The keyword stripping step is aided by an optional automatized step that consists in extracting the context in which a word appears to verify its relevance.

First, we manually collected a representative sample of literature, books, and articles, relating to tool use and tool making among primates, which we refer to as the ‘corpus’. The corpus is representative in the sense that it provided sufficient vocabulary to start the process of describing behavioural processes. We set out to compare primate tool use. Because of the comparatively sparse data on organic tool use amongst hominins, we focused here specifically on percussive tool use with lithics. The corpus contains 75 documents whose publication dates range from 1957 to 2021, with most of the publications being from the 2000’s (average: 2004, median: 2009). 54 documents (72%) primarily concern primatology. Nineteen (25%) documents concern formal methods for describing and producing models of animal tool use, such as (Hayashi 2015). Articles in which behaviours and cultures are compared (e.g., Leca, Gunst, and Huffman 2007; Whiten 2011) are particularly valuable. Generally, these comparisons of cohorts of the same species, such as the common chimpanzee *Pan troglodytes*, include a more detailed behavioural and material description compared to inter-species comparisons: thus, 17 (22%) documents are related to the concept of ‘culture’ and cross-community variations.

We then extracted detailed tool use descriptions from the corpus by taking special attention to the context. Using the Rapid Automatic Keyword Extraction (RAKE) algorithm (Rose et al. 2010) we first obtained a raw list of words and phrases, i.e., expressions of several words such a ‘human evolution’ which appears in 39 (52%) documents in the corpus and ‘wild chimpanzees’ which appears in 43 (58%) documents. This raw list was then manually stripped of irrelevant data such as *stop words* that carry little information, for example ‘be,’ ‘related,’ ‘is,’ ‘he’, or words that are not related to tool use and tool making. In natural language processing and linguistics, the *context* in which a word appears in a text can be defined as the words or the characters that appear around the word. The *left context* is the context that appears *before* the word; the *right context* is the context that appears *after* the word. To help verify the relevance of the resulting list of keywords and discover relevant words that might have been missed by the automatic keyword-extracting algorithm, we thus also extracted the context in which the keywords appeared. In Figure 5, we illustrate how the context can help both verify the relevance of a concept and the discovery of new concepts. We give a detailed description of the main sets of concepts we have added into the ontology in Section 3.

**Figure 5:**
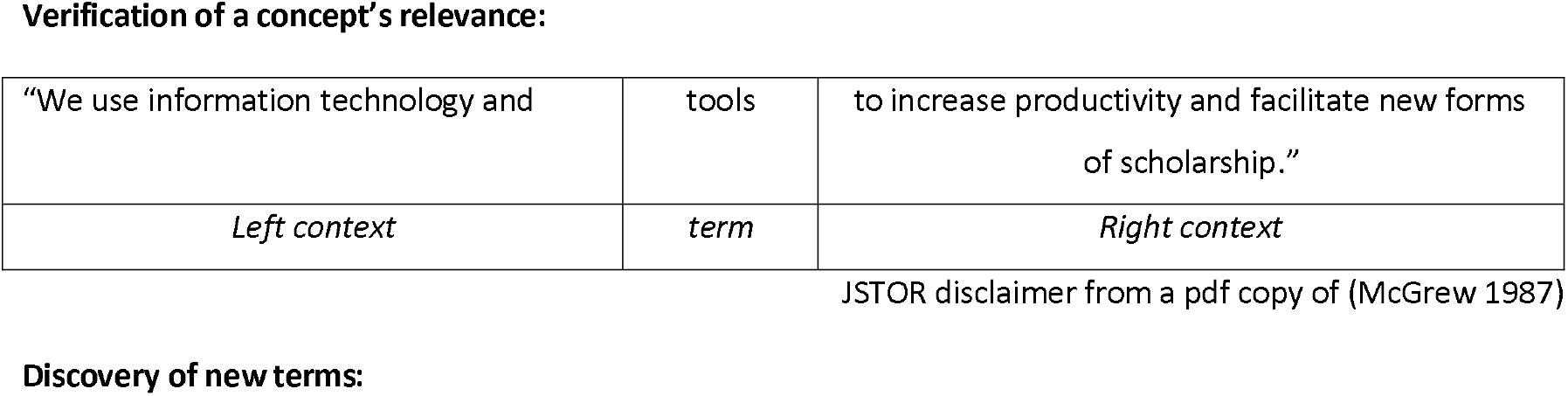
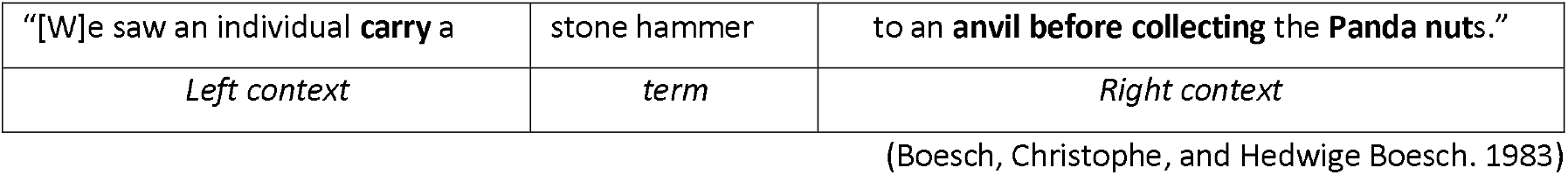
Examples of results for the search of contexts across the corpus. Top: a seemingly relevant term ‘tools,’ extracted automatically, appears in a context that is not relevant (*noise*). Bottom: from the automatically extracted keyword ‘stone hammer,’ we gain knowledge of related concepts, indicated in boldface that appear in the contexts, such as ‘anvil’ and ‘Panda [nut],’ but also the concept of carrying and collecting something, as well as the concept of sequential actions.

Besides concepts, we also extracted and manually selected *relations between concepts*. A relation, in mathematics and information science, is a set of couples that link elements together: for example, the relation *olderThan* can be defined as the set of couples of things (A,B) where the thing A is older than the thing B. LOM3-2011-I16-3, a unifacial core-tool from Lomekwi and estimated to be 3.30 million years old (Lewis and Harmand 2016), is older than the LA-91-W-47-9 core-tool from Lokalalei, estimated to be 2.30 million years old. We can express the relation between those two items as:

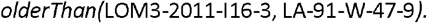

As an illustration of the process of extracting concepts and relations, consider the following example of the alpha-male ‘BT’ honey-dipping in a tree hole (Watts 2008, page 90):

> *“– July 16, 2002: Four adult males found a beehive in a hole in a fallen tree trunk. (*…) ***α-male BT (***…***) inserted a stick about 30 cm long and 1.5 cm in diameter into the hole. He thrust it back and forth and from side to side vigorously, then withdrew it and ate a small amount of honey that adhered to it. He repeated this several times[.]”***

To capture this concrete ethological example of manipulation entirely into our ontology, we will require the introduction of concepts and relations for the chimpanzee ‘BT’, his actions, the targets of those actions, but also for the way those actions are linked together. For example, we want to indicate clearly which action come after the other.

We may first focus on the concepts related to the agent. That is, the individual performing the action: *Male, Adult, Chimpanzee, Alpha*, which have taxonomies associated to them. For example, the concept of Chimpanzee is stored within a phylogenetic taxonomy of primates in our ontology. Similarly, geographical concepts such as *Ngogo, KibaleNationalPark, Uganda* are associated with a simple geographical taxonomy. Relations such as *age, sex, location* provide additional structure to relate the relevant concepts together and further describe features of the agents at play.

The most important relations that we extracted are the ones related to behavioural descriptions. Concepts such as *Tool, Stick, OrganicTool* allow us to describe the tool being used. We define specific concepts to describe the way a tool is being used: *Thrust* and *ThrustBackAndForth, ThrustFromSideToSide, Pull*, but also *EatHoney*, which is the end goal of the overall behaviour. These concepts are high-level for now, but they can easily be given alternative definitions depending on the needs of the researcher. They can also be defined from more fundamental motions that can be reused across a wide range of different situations. For example, the concept *EatHoney* that corresponds to the notion of eating honey can be instead made up of the concepts *Eat* and *Honey* which allows us to reuse the concept *Eat* in a different context. Ant-dipping is another context where the chimpanzee will instead dip a stick inside a hive to fish for ants (McGrew 1974). These concepts were inspired by ethogram literature (Nishida et al. 1999) and the specific vocabulary from the raw descriptions themselves. The target of the action also has its own concepts and taxonomy: *Honey, Beehive, Hole, TreeTrunk, Tree*. Finally, concepts that correspond to describing behaviour tie everything together, such as the most general and abstract concept of *BehaviouralInstance*. These raw descriptions are instances of the more specific concept of *Bout*, which refers to the description of the behaviour itself, and *SubBout* into which a *Bout* decomposes. Here, the *Bout* may be decomposed into several sub-bouts for BT’s actions of inserting the stick into the hole, thrusting it, withdrawing it, and eating honey from it. How to decompose actions and how to arrange them is a challenge (Lashley 1951, Rosenbaum et al. 2007). For this exploratory work, we simply choose to follow the choices of the textual descriptions’ original authors as closely as possible. If necessary, the decomposition of actions can be made more precise or changed by adding or modifying concepts in the ontology, which is easy. We defined specific relations to tie sub-bouts together in a directed graph manner reminiscent of Petri nets or automata, which can be seen as directed graphs that model discrete temporal events or sequences of events. In Figure 7, Section 3, we show graphically how this specific instance of behaviour can be stored into the ontology and how similar it is to automata where one may go from one node to another following arrows.

### 2.2. Adding structure to the vocabulary list: tool and tool function taxonomy

Now that we have a list of vocabulary relevant to tool use and tool making, we need to add structure to them. This is important for later automatic reasoning and data analysis. For example, thanks to a taxonomy for primates as in Figure 1, a machine can easily ‘know’ that a wild bearded capuchin is a capuchin because of the conceptual, semantic relation *WildBeardedCapuchin isA Capuchin*.

We used an automated process to build part of the taxonomy from unstructured descriptions of tool behaviour and tools from (Whiten et al. 1999). Using Formal Concept Analysis (Davey and Priestley 2002, Bendaoud, Toussaint, and Napoli 2007), we analysed these descriptions of the tools, which includes their purpose and the different stone materials they are made of. This formal mathematical method gave us a taxonomy by ordering tools according to their shared features. For example, tools that are made from the same materials are classified under the same category. This gave us many subsumption relations between concepts that can be reused later (Codocedo and Napoli 2015).

Formal concept analysis starts off by describing tools and their features in a so-called binary table. Here we classify tools according to the behaviour associated with them. Each row in the corresponding binary table is a different type of tool, and each column corresponds to broad categories of behaviours. An ‘X’ at the intersection between row and column indicates that there is documented use of the corresponding tool for the corresponding behaviour.

In Table 1, we extracted descriptions of tools and their functions in (Whiten et al. 1999).

**Table 1:** Some tool functions among chimpanzees related by behaviours. X indicates the use of a tool during a specific behaviour. PO: pounding behaviour, such as using a hammer to crack open a nut; FI: fishing behaviour, such as termite-fishing with a leaf midrib; PR: probing, such as picking bone marrow out or flicking bees to access honey; FO: making use of force, such as using a stick to enlarge the entrance to an insect hive; CO: comfort-related behaviour, such as using large leaves as seat or whisking away flies with a leafy stick; EV: ‘miscellaneous exploitation of vegetation properties’, such as using objects to self-tickle; EL: exploiting leaf properties, such as squashing an ectoparasite or inspecting it once it is on a leaf; GR: grooming, which also includes squashing ectoparasites; and AS: attention-seeking behaviour, such as noisy branch slapping, knocking knuckles, or noisily pulling stems (data from Whiten et al 1999).

| Tools | Behaviours |  |  |  |  |  |  |  |  |
| --- | --- | --- | --- | --- | --- | --- | --- | --- | --- |
|  | PO | FI | PR | FO | CO | EV | EL | GR | AS |
| **branch** |  |  |  | x |  | x |  |  | x |
| **grass** |  | x | x |  |  |  |  |  | x |
| **large_branch** |  |  |  |  |  |  |  |  | x |
| **leaf_stem** |  |  |  |  |  |  |  | x | x |
| **stick** | x |  | x |  |  |  |  |  |  |
| **stone** | x |  |  |  |  | x |  |  |  |
| **stone_anvil** | x |  |  |  |  |  |  |  |  |
| **stone_hammer** | x |  |  |  |  |  |  |  |  |

This table allows us to mathematically extract concepts, that are partially ordered according to the functions they share or, dually, the behaviours a certain set of functions share. For example, Table 1 defines the concept of tools that can be used for both grooming (GR) and attention seeking (AS):

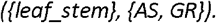

In Figure 6, we represent the concepts of Table 1 and the partial order they are in as a line diagram, also called a *concept lattice*. In the diagram every node corresponds to a concept such as ({leaf_stem}, {AS, GR}), that we add to the ontology. Furthermore, the nodes are ordered by set inclusion of their intent and extent: in two linked nodes, the one situated lower in the diagram represents a more general concept. This means that the concept lattice is a taxonomy of tools and functions, that we can add to our ontology.

**Figure 6:**
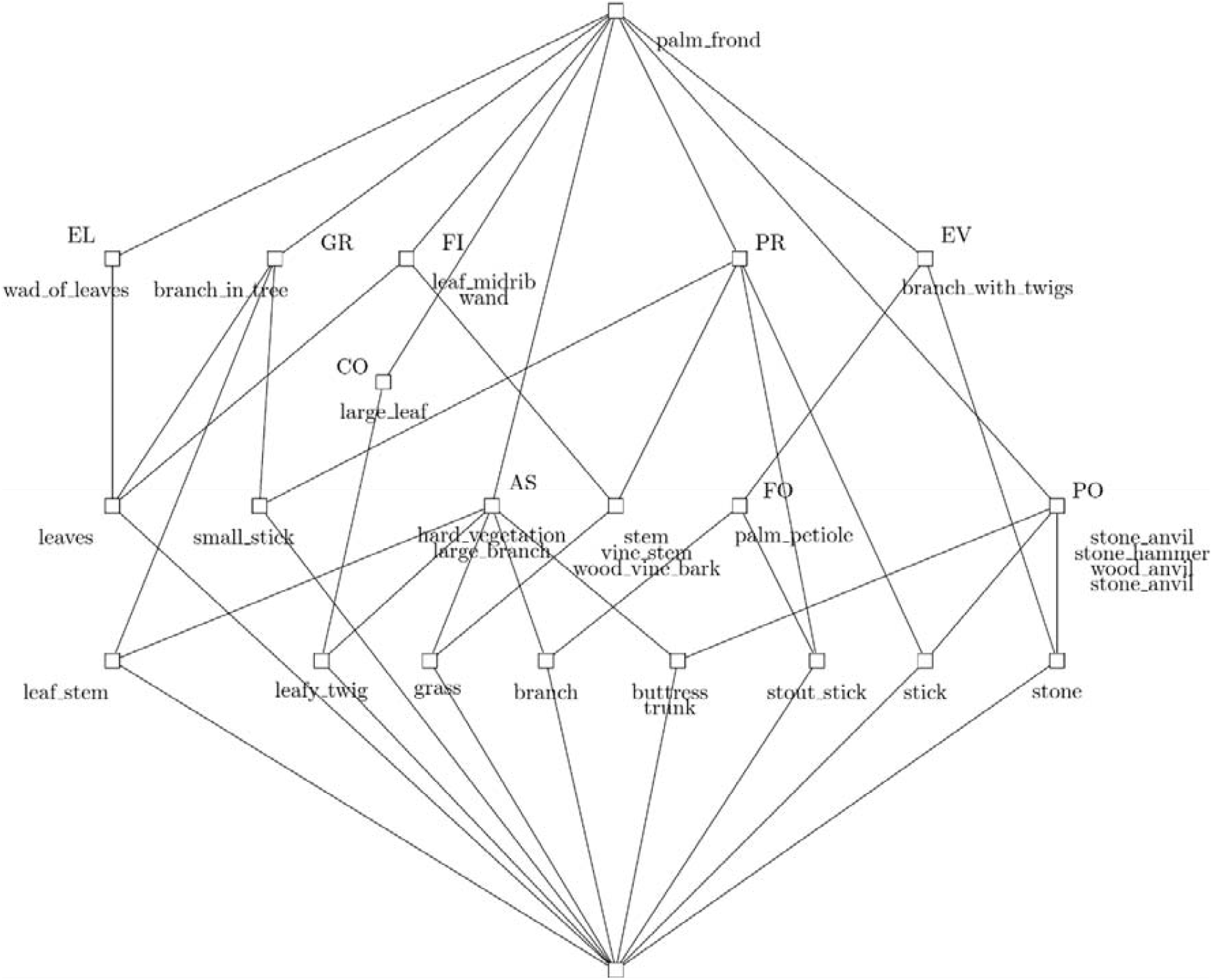
Concept lattice as defined by Table 1. Concepts are represented here as nodes. Generated with *GraphViz* and *Concept* modules for Python (Bank [2014] 2021).

#### Building the ontology

Now that we have our concepts and hierarchical relations between them, we can build our ontology so that it can be read by a machine. To formally define the concepts and the relations between them, we used OWL (Web Ontology Language) (McGuinness, Van Harmelen, et al. 2004). OWL allows us to indicate more precise information about the concepts and relations. For example, using OWL we can define two concepts as being disjoint, which means that there cannot be any instance of both. This is the case of the concepts *Macaque* and *Chimpanzee*, and of the concepts *Stone* and *LeafMidrib*. However, *Tool* and *Stone* are not disjoint, because there exist tools made of stone. Using OWL, we can also indicate the domain and codomain of some relations^1^. For example, the relation *describedInText* links the concepts *BehaviouralInstance* and *Text*, which means that an instance of *BehaviouralInstance* can be linked, through *describedInText*, to an instance of Text. This expresses in a machine-readable way the fact that a certain behavioural instance is described in a certain text, as with our running example of the honey-dipping chimpanzee.

To construct the ontology, we used the Protégé software (Noy et al. 2003, Musen 2015), starting from a part of the reference ontology SOMA. Because SOMA (Socio-physical Model of Activities) is used to provide autonomous robots with knowledge regarding everyday activities and human common sense, it is an excellent basis for describing the concrete concepts and objects including tool material and physical anatomical characteristics. Also, SOMA provides us a basis for describing abstract concepts related to tool manipulation, such as tool orientation and body posture.

In Figure 7, we demonstrate the thoroughness and representation power of the ontology by representing our running example of the honey-dipping chimpanzee BT. Note that for clarity’s sake the graph does not contain all hierarchical relations: for example, the subsumptions that follow that BT is an *Agent-Generic* were left out.

**Figure 7:**
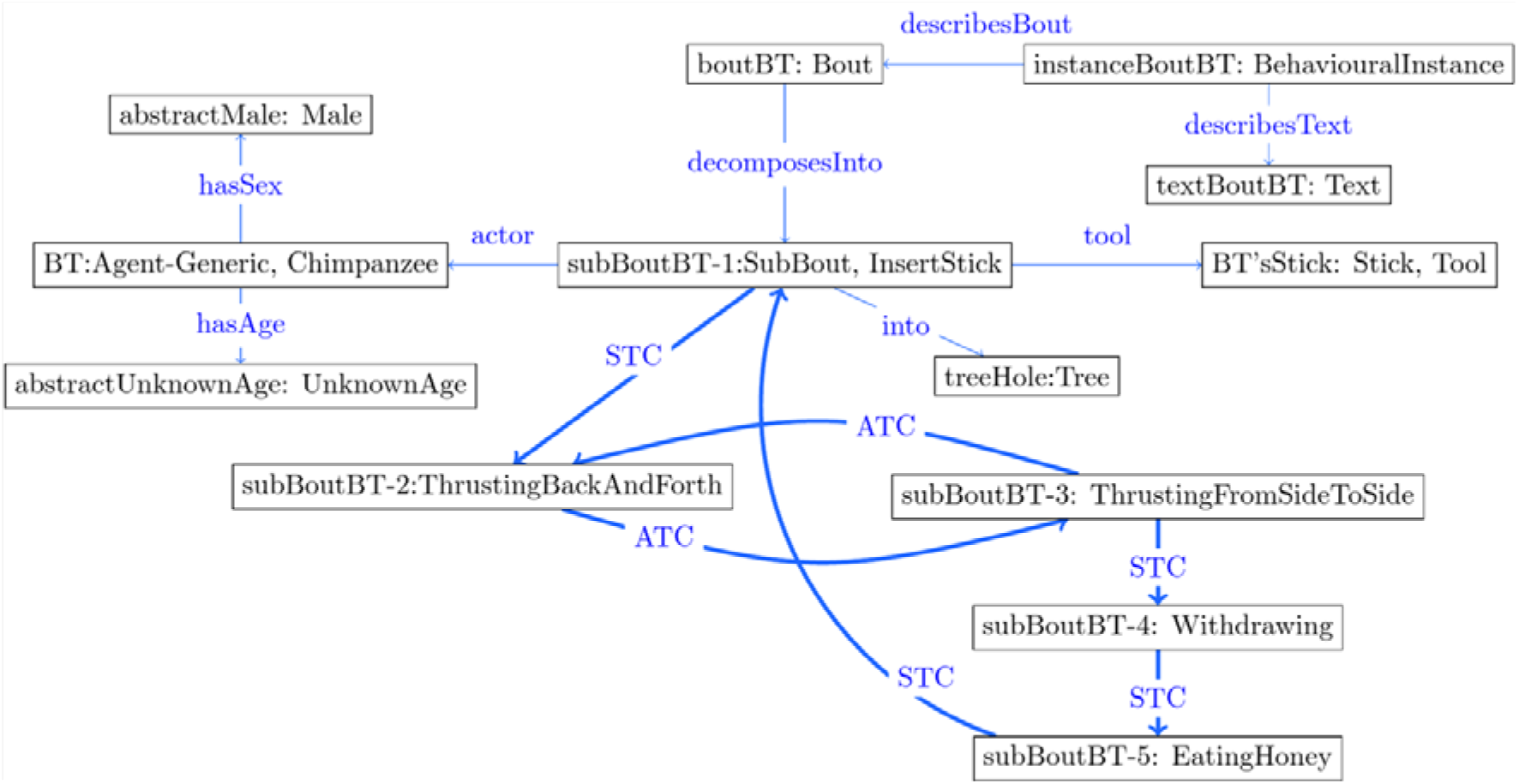
Partial graphical representation of a sequence of actions that describe a chimpanzee, BT, who dips a stick inside of a beehive to extract and eat the honey. Instances are given in boxes following the object-oriented notation ‘nameOfInstance:Type’: thus ‘BT:Chimpanzee’ indicates that ‘BT’ is an instance of the concept *Chimpanzee*. Arrows between instances denote the relations between instances. STC and ATC stand for the relations *sequentialTemporalConjunction* and *ambiguousTemporalConjunction* respectively, indicated by thicker arrows in the figure. The separation of actions is as follows: (1) BT inserts a stick inside of a hole; (2) BT thrusts the stick back and forth and (3) from side to side; (4) BT withdraws the stick; (5) BT eats honey; (6) BT repeats the process. For clarity’s sake, only the first instance of *SubBout, subBoutBT-1*, has been entirely related to *BT* and the instances related to him. For the same reason other information, such as where exactly the bout happened, were left out.

Finally, we built a new dataset and added explicit descriptions of primate behaviour related to tool use and tool making. We curated 365 articles, books, and communications, from 1887 to 2021 (mean publication date: 1995) of which 65 (17,8%) were relevant in the sense that they contained explicit descriptions of behaviour. Publications were deemed irrelevant to our purposes if they contained statistical interpretation on primate tool use or tool use in captive primates. For example, (Hobaiter and Byrne 2011) describes a repertoire of chimpanzee gestures which is particularly relevant as a solid source of concepts such as *LeafClipping or ObjectShaking*, but does not describe individual-specific use of gesture as the running example of honey-fishing. Rather, they perform statistical analyses and compare groups of individuals to guarantee the significance of their gestures.

This dataset was created with the help of experts in primatology and anthropology. Note that the corpus from which we extracted our starting vocabulary and this new dataset share part of their entries. This new dataset is much larger and specific to primate behaviour: extracting a starting vocabulary from this new dataset would thus have yielded too much vocabulary to curate and denoise. From these articles, we extracted more than 200 instances of behaviour involving tools. The majority (64%) involve chimpanzees, followed by capuchins (19%), gorillas, macaques, baboons, orangutans, red colobus monkeys, spider monkeys and red howler monkeys. Most tools are organic (90%), and includes leaves, twigs, wooden sticks, hammers, or anvils. Stone tools (5%) are exclusively used in the context of hammering, i.e., as hammer or anvil. The descriptions range from the simple description of a single action, e.g., using a stick to clean one’s nose, to a sequence of several actions, such as ant or honey-dipping. The 200 instances of behaviour were described entirely, meaning they have been represented as sequences of behaviours in the ontology, including our running example of BT’s honey-dipping.

## 3. Results

We now describe the contents of the ROCA ontology after the building process explicated in Section 2. We describe the concepts and relations of the ontology. We describe the instances of behavioural descriptions of non-human primate tool use and tool making that we have incorporated in our ontology. We also show how to obtain additional knowledge from the ontology and what type of knowledge by performing automated queries and analysis on its data.

### 3.1 Description of the contents of the ROCA ontology

Our ROCA ontology contains more than 900 concepts, including more than 700 new concepts compared to the starting SOMA ontology (https://rocaontology.github.io/). In this section we describe the main sets of concepts, the main sets of relations, and examples of descriptions included in the ontology. To indicate clearly which concepts come from SOMA, we use the syntax ‘SOMA:concept’.

#### Concepts to describe physical movements

Most of the movements we newly included correspond to sub-concepts of ‘SOMA:Translocation’. That is, moving something from one place to the other, like, for example, *LiftingAnObject* and *LoweringAnObject*. This also includes *EdgeHammering, FaceHammering, PointHammering*, which are all sub-concepts of Hammering itself and depend on the area of the object that is being struck. More concepts allow us to describe the posture of the agent such as Posture-Configuration, which include *Rising, Sitting, Standing, StandingBipedal, StandingTripedal*. Agents can also grasp a tool differently, which is described with concepts such as *HoldingWithHand, BothHands, GraspingSomething* with a *PowerGrasp* or a *PrecisionGrasp* with a *PalmarPinch*. Most of those movements were extracted from (Gumert, Kluck, and Malaivijitnond 2009; Tan et al. 2015), as well as works with the specific aim to build general (Pastra and Aloimonos 2012) and primate-centered (Hayashi 2015) grammars of actions. The productions (sentences) in these grammars are sequences of events encoded with shorthand symbols, much like in ethology and behavioural observation in the wild, such as the sequence ‘PHT’ for ‘pick, hold, throw.’ These sequences are reliant on an additional lexicon to define what the symbols stand for and are usually heavily dependent on the context: what was picked, held, thrown is supposed to be clear from the context and does not necessarily appear within the grammar. In our case, we add onto this type of representations the expressive power of directed graphs: we can make the target of the tool use explicit in the representation.

#### Abstract concepts to relate tools, targets, and agents

*SOMA:Situation, SOMA:Event*, and their types such as *SOMA:SeparationEvent, CuttingEvent*, and *Cracking* (in the case of nut-cracking) allow us to relate tools, targets, and agents via the appropriate relations, as in Figures 2 and 7. For the descriptions of an event of tool use that does not require a sequential description of movements, we have chosen the notion of *B*out as central: recall that a ‘bout’ is a period of activity during which a tool is used. In observatory papers, it often corresponds to a period that starts from the agent picking up the tool and ends with the tool being discarded.

Two events with a certain length in time can be compared using so-called ‘interval algebra’. For example, climbing a tree takes a certain amount of time, and so does picking up a fruit in the tree. An interval algebra allows researcher to precisely indicate that climbing the tree comes before picking up a fruit, or perhaps that dangling from a branch occurs while picking up the fruit. For our purposes we used a simplified version of Allen’s time interval algebra (Allen 1983). We use the relation *parallelTemporalConjunction* for events that occur in parallel and the relation *sequentialTemporalConjunction* for events that occur one after the other (cf. Pastra and Aloimonos 2012). With these we can describe and recover a temporal order, that is, an order that depends on time, of the actions performed during the bout, such as picking up a tool, *then* using it, *then* discarding it. We also include the relation *ambiguousTemporalConjunction*, when there is not enough information within the description to decide whether events are related sequentially or in parallel. We can thus avoid the more complex grammatical requirements described in (Pastra and Aloimonos 2012), such as long-distance dependencies, by making use of the graphical nature of our representations and the sharing of sub-graphs. For example, when a procedure calls for the repeated use of the same stone hammer, we can always refer to this same hammer instance directly within the structure of the representation. Notice the similarity of this conceptualization with directed graph-based representations such as Petri nets (Murata 1989) and finite state machines (Savage 1997).

#### Concepts to describe the agents and tools

We created concepts to describe the agents, the entities that perform (act) an action. These concepts describe the tool-using agents in the ontology and were obtained from the literature, for example *Bonobo, Capuchin, Chimpanzee, Orang-Utan, Macaque*. Some of these concepts include additional information if necessary: *BurmeseLongTailedMacaque, JapaneseMacaque*. The latter is made more precise with corresponding scientific names as *MacacaFuscataFuscata or MacacaFuscataYakui*. Hominids descriptions also benefit from pre-existing classifications such as *HomoErectus, HomoErgaster, Denisovan, Neanderthal*. Agent specific information is captured by concepts such as *Age, Adolescent, Adult, Juvenile, Sex, Female, Male*. Other concepts are used to describe the target of tool use and the tool itself. The physico-chemical characteristics of tools can be described, such as its Material: *Basalt, Sandstone, Diorite, Granite, Quartz* (Tan et al. 2015). The characteristics of the target can for example be described as *Nut, NutShell, NutHardness, OilPalmNut, CoulaNut, TucumNut, ManihotNut*, but also *Bone, BoneMarrow, Seafood, Seashell*. Finally, some other concepts allow us to add additional information to the tool use event itself. These concepts express geographical information to describe areas where observations were made, areas of tool procurement, and areas of tool excavation. For example, *OlduvaiGorge* in *Tanzania, EastAfrica*, and Bossou in WestAfrica where many wild chimpanzee studies have been carried out. Some concepts also express temporal information on *Season*, with *DrySeason* and *WetSeason* for *TropicalSeason*, and concepts related to *Weather*, but also specific *StoneIndustry* such as *Acheulean, Aurignacian, Mousterian*, and *Oldowan*.

### 3.2. Knowledge acquisition: Queries and statistical analysis

Now that our ROCA ontology for tool use and tool making is ready with concepts, relations, and instances, we show how this data can be used to derive new knowledge, and how the specificities of the ontology make this knowledge discovery easier.

The reasoning over the ontology allows us to obtain automatically new knowledge about the domain of tool use. Querying the ontology is one way of reasoning. Direct queries can be performed from a standard automated reasoner, such as Fact++ (Tsarkov and Horrocks 2006), that is directly available in the Protégé ontology editor software. These queries return sets of concepts or sets of instances from the ontology. Here we elaborate using several example questions. The first being:

> *What different types of hammering are there?*

From the ontology, we obtain the specific concepts pertaining to “hammering”:

> *Hammering itself, EdgeHammering, BimanualAsymmetricalEdgeHammering*,
>
> *BimanualSymmetricalEdgeHammering, UnattachedEdgeHammering*,
>
> *UnimanualEdgeHammering, FaceHammering, BimanualAsymmetricalFaceHammering*,
>
> *BimanualAsymmetricalFlipFaceHammering, BimanualFulcrumFaceHammering*,
>
> *BimanualStandFaceHammering, BimanualSymmetricalFaceHammering*,
>
> *UnimanualClapFaceHammering, UnimanualFaceHammering*,
>
> *UnimanualFulcrumFaceHammering, UnimanualShellFaceHammering, PointHammering*,
>
> *BimanualPointHammering, UnimanualJabPointHammering*,
>
> *UnimanualPointHammering, UnimanualShellPointHammering*,
>
> and *UnimanualSupportedPointHammering*.

The results of this query are useful in at least three ways. First, it allows researchers working on the description of behaviour to immediately obtain an ethogram to describe an instance of hammering. Second, the elements of this ethogram are already ordered hierarchically. Thus, a researcher can decrease the degree of complexity in their ethogram by choosing the more general concepts EdgeHammering, PointHammering, and FaceHammering, or focus on the Unimanual nature of the hammering depending on the focus of the analysis. Third, with the addition of concrete instances of behaviour, a researcher can obtain statistical data for each sub-concept. In our case, we identified that, given our own corpus of descriptions of instances and the concepts we have added in our ontology, hammering associated with stone shows the most variety when compared with hammering associated with wood; these concepts only appear in the context of stone tool use, whereas wood tool hammering is never described using these precise concepts. This may indicate either a bias in our corpus towards tool descriptions or suggest that wood hammering has not been described with the same degree of precision as stone hammering, or that wood hammering is intrinsically less complex, in that sense, than stone hammering.

In another example query, we explore female primate tool use. The ontology contains instances of female primates. For example, from (Lawick-Goodall, Lawick, and Packer 1973): “One three-year-old female, however, picked up a fairly large stone with which she repeatedly and forcefully rubbed at her muzzle in an attempt to remove the dried juice.” We can use these instances to answer questions like:

> *How many cases of female tool use are there in the ontology, and which individuals are they?*

The ROCA ontology contains 126 individuals that are female primates involved in cases of tool use or tool making (Figure 8).

**Figure 8:**
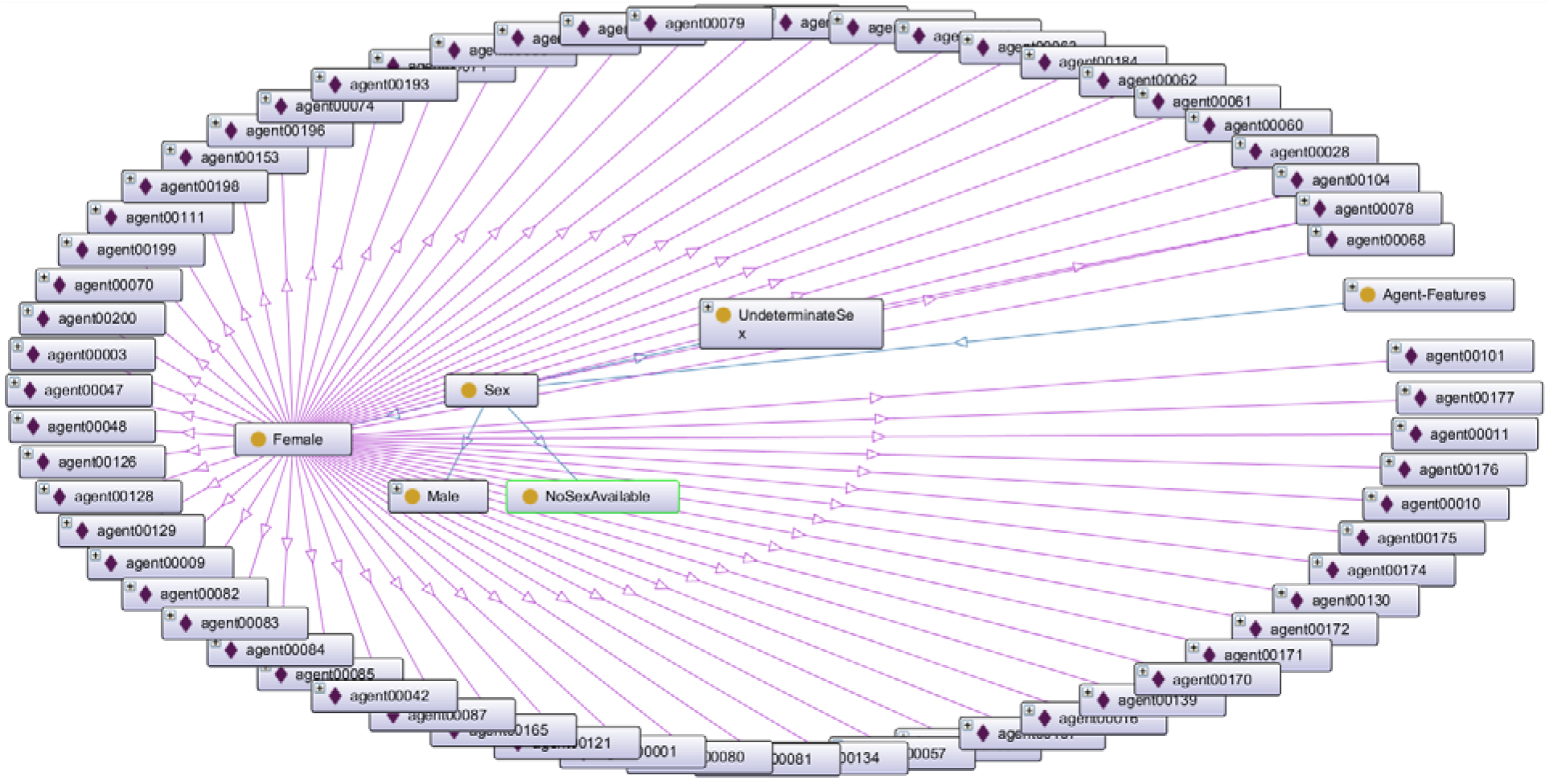
An ‘ontograf’ view of part of the ROCA ontology in the Protégé software. Instances are indicated as boxes with a lozenge, and concepts are indicated as boxes with a circle. All the instances displayed are instances of the *Female* concept.

By representing the sequences of actions explicitly, we can also answer questions specifically related to those sequences and verify hypotheses related to tool use. For example:

*Can we conclude whether an individual is male or female based on their tool use behaviour?*

To answer this question, we first study the lexical similarity of instances of behaviour only in terms of the actions that appear in the sequences, and next we analyse the results using a clustering algorithm to hopefully extract a pattern. The lexical similarity between two instances of behaviour is a number between 0 and 1, defined as the number of actions they share divided by the number of actions described in both instances. For example, the lexical similarity between *“Picking, Biting, Crunching, Standing, Picking”* and *“Coming, Shaking, Biting”* is 1/6. The higher the number, the more similarity between the two groups in the comparison. In Figure 9, we represent a similarity matrix between all 200 instances in the ontology.

**Figure 9:**
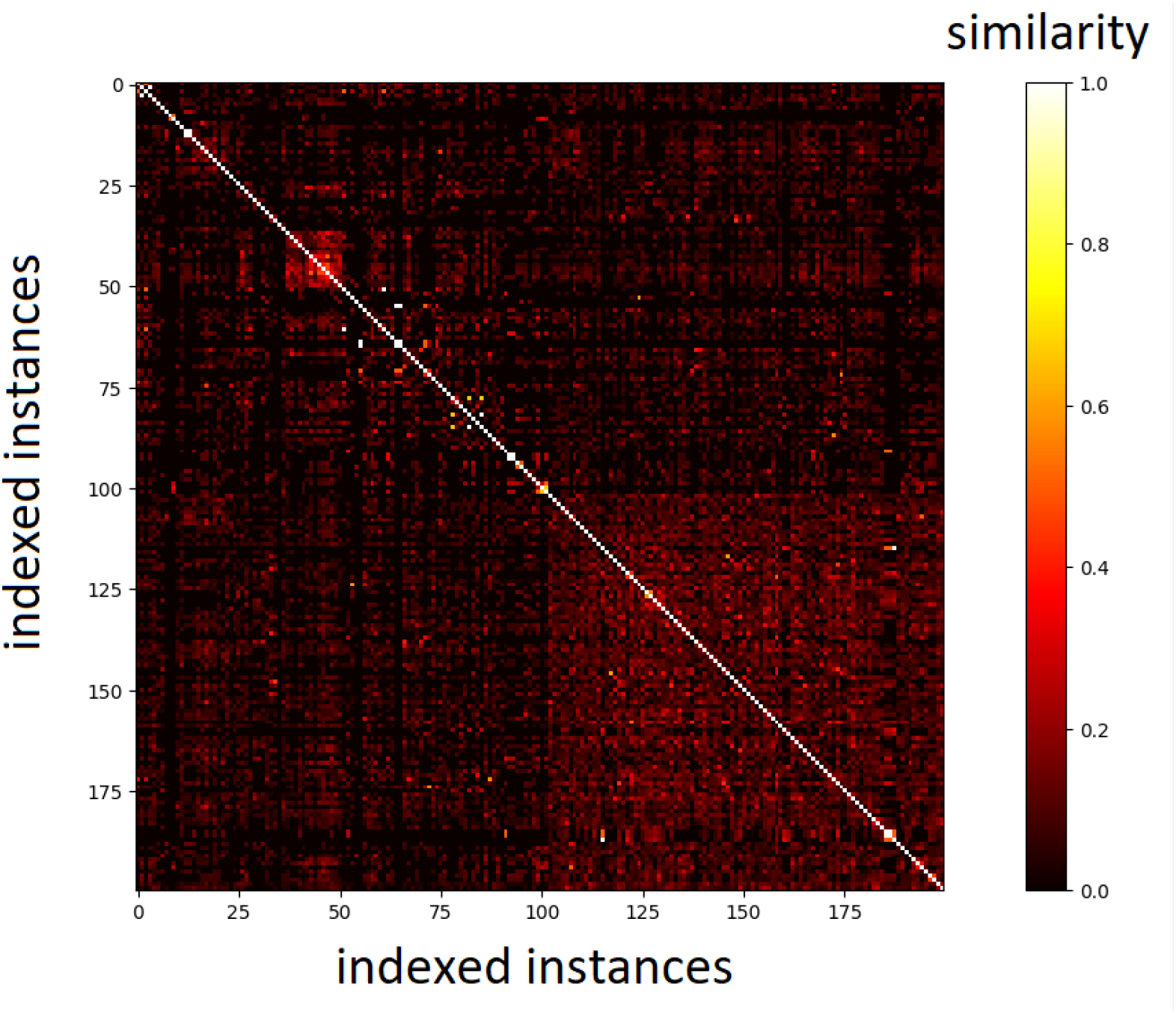
Sorted similarity matrix between instances of behaviour. Each dot corresponds to the similarity between two instances, which are indexed in both axes. The brighter the dot, the higher the similarity. Large clusters appear as large rectangles.

To identify a pattern in tool use and gender we used the clustering algorithm K-means (Steinhaus 1957, MacQueen 1967). The K-means allows us to classify the instances into K=2 groups according to their similarity. The K-means is represented by precision number, which is defined as the ratio between true positives and the sum of all detected values. A precision close to 0.5 means that 50% of positive group identification were correct. The hypothesis is that the gender of the tool user has an impact on the similarity of sequences of behaviour. If descriptions of female tool users are more like each other (more similar) than descriptions of male tool users, then we will recover a male/female split by a similarity number far from 0.5. We ran the K-means Python implementation from scikit (Pedregosa et al. 2011) on the similarity matrix and obtained a precision number of 0.48. This means that using the lexical measure on the descriptions of behaviours to recover a male/female split is not better than choosing which instance corresponds to a male or a female individual at random.

Thus, the description of sequence of behaviours alone cannot help recovering the sex (male or female) of the tool user. One interpretation is that there is no significant lexical difference between *descriptions* of sequences of behaviour of male and female individuals, and thus that there are no apparent differences between male and female tool use in terms of the words being used to characterize their actions, inside of our corpus.

## 4. Discussion and Conclusion

The ROCA ontology (https://rocaontology.github.io/) is a newly created and freely available tool that can **handle knowledge obtained from the literature, by making the data uniform, unified, acentric, dynamic, and human-readable, to allow for deeper analysis of primate tool use**. The data-mining tools we created to populate ROCA ontology is also freely available from the ROCA platform.

### 4.1. Qualitative evaluation of the ontology

We set out to study if ontologies can provide a unifying framework to describe and reason over tool use. Here we assess the ROCA ontology from our exploratory work we can conclude that ontologies provide a **uniform, unified, acentric, dynamic, and human-readable way to handle knowledge obtained from literature**. Thus, overcoming current research problems of chaînes opératoires. Also, ontologies allow us to move beyond the current state-of-the-art, and facilitate new types of knowledge acquisition.

#### Uniform and unified data

Instances and vocabulary have been aggregated to a singular place, the ROCA ontology. Instances use only the concepts that are defined within the ontology, no matter their source. This apparent constraint forces the data to be uniform semantically. Structurally, one can define guidelines to describe behaviour in the same manner. The benefit is twofold: it helps specialists record and analyse their own data, and it helps compare any instance of behaviour with any other. This can be valuable for archaeologists seeking to compare data chronologically or geographically, to highlight cultural differences for example.

All our data has been aggregated from many different sources and regrouped to a singular place, which makes data mining and subsequent analysis easier across all those sources. The ontology is thus more than a way to unify data and analyses, it also functions as a database. More work is needed to explore how to meaningfully include sources than texts, such as pictures, videos, physiological measures, and 3D models. The first three can be considered closer to direct observation without the degree of interpretation brought by a researcher’s analysis and textual description which might introduce unwanted biases.

#### Dynamism

Data can be continuously added and modified to the ontology, whether it concerns new concepts, concept definitions, relations between concepts, and instances of, e.g., tool use. It is straightforward to include data related to tool use by other non-primate animals, such as crows, insects, or cuttlefishes, or instances of experimental research in behavioural biology and archaeology, like reconstituting toolmaking by knapping stones. This is highlighted by the process of creating the ontology itself: we started with a vocabulary, which we gave structure to; later, when we added instances of tool use by chimpanzees, we simply added more concepts (see Section 3.1). One specific example is the diversity of motions we identified in the second survey. We created new concepts from the raw text because the results from the first vocabulary survey were not specific enough anymore. We added, for example, *Stomping, Dunking* or *DunkingInWater, BringingOut*, and *Thrusting*.

Ontologies are can easily be augmented and maintained to capture more types of tool use and manipulations, of tool building, as well as more types of agents, for example different hominin species, or other animals that are known tool users such as corvids (Hunt 1996) and octopuses (Finn, Tregenza, and Norman 2009).

#### Acentricity

Compared to strictly hierarchical taxonomies and textual descriptions, ontologies are acentric and can be used by researchers with different interests and questions. One may choose on which concept or agent to focus such as the tools, the tool users, the tool materials, as highlighted by the query results in Section 3.2. In our second example we can ‘filter’ the instances of the ontology to only consider *tools that are made of Quartzite* and everything that is related to it. We can also filter the ontology to only consider, *female chimpanzees involved in hammering*. The focus for both is different: where we would need several databases or several taxonomies - one for tools, and one for chimpanzees -, the ontology contains both and the centre is chosen by the user.

#### Human-readability and variability of representations

Because levels of hierarchy can be left out or included the level of ‘data noise’ can be manipulated, making data easier to handle for both humans (readability) and machines (computational complexity). Consider for example Figure 10, that represents a generic instance of oyster cracking, but on which there are much less concepts involved than in Figure 7. Both these figures are centred around the concept of Bout. One can control the depth of description by allowing a controlled number of concepts to appear or by restricting the types of concepts that appear, such as only showing the tool and the agent involved. In practice this can be done by restricting the number and the type of nodes: in Figure 10, all the nodes are separated from Bout by at most a single node, which helps keep the graph small and readable.

**Figure 10:**
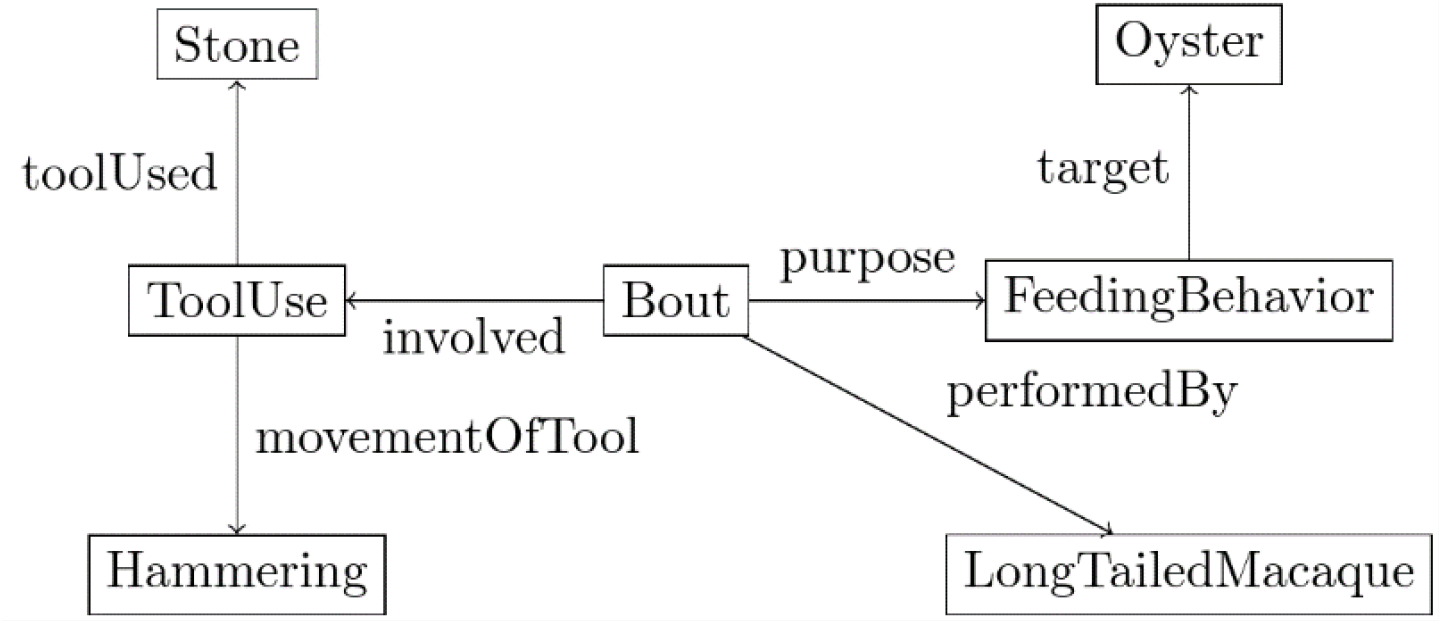
Simple directed graph representation, or concept graph, of an instance of oyster cracking.

#### Knowledge acquisition

Ontologies also allow for knowledge acquisition via queries which we illustrated above. They allow us to derive new knowledge via automated reasoning, like the lexical similarity between instances and not only via statistical analysis, like K-means.

#### Quality of the ontology and representativity of the data

The ROCA ontology is the proof of concept that demonstrates the power of ontologies in behavioural research. However, we note several points for improvement. First, the quality of the descriptions obtained from text is required. A manual assessment and verification of the quality of the transcription ‘text to instance in the ontology’ is needed to ensure that the instances correspond exactly to what the textual behavioural descriptions tried to convey. Second, in the compilation we focused on the existence of a behaviour, and not the statistical significance of it. Because we were interested in describing instances of tool use, we recorded for example, instances of nut-cracking with rocks. As such we were not interested in having many different records of nut-cracking to confirm trends statistically. It was enough for us to know that at least one primate used a tool in a certain context. With only 200 instances of behaviour, of which 5% involve stone tools only, the ontology is not large enough to infer reliable trends regarding stone tool use and tool making. Third, the concepts and relations that are used in the ontology need to be defined precisely to ensure that the vocabulary relates to consistent concepts across representations. An alternative is to define additional concepts or aliases for things that are not synonymous in different fields. For example, *Cracking or Breaking* are near-synonymous, and should appear as such in the ontology by way of a (semantic) relation. This is like the semantic organization of concepts in the lexical database for English, WordNet (Miller et al. 1990) in which words are grouped according to their meaning. On the other hand, two concepts might be dissimilar despite having the same name. For example, (Gosden and Head 1994) describe the ambiguity of the term ‘landscape’. To solve this ambiguity by way of an ontology, we would define a concept for *TransportedLandscape* to refer to the “fact that most of the useful plants and animals (…) were taken there by people” (Kirch 1984, 135, p. 9), *TransformedLandscape* to express the modification of flora and fauna by prehistoric inhabitants (Head 1994), *VisualLandscape* to focus on the signs and marks on the land (Tacon 1994), *PhysicalLandscape, SocialLandscape* (Pickering 1994), and as many remaining concepts as we would need. We would then leave it to the researcher to use the conceptualization of ‘landscape’ that is the most relevant for their work.

### 4.2. Assessment of the Ontology

#### Complex structural queries

In Section 3.2, we queried the ROCA ontology to derive new knowledge. We obtained results that were sets of concepts (‘What are the concepts associated with hammering?’) or sets of instances (‘Who are the female tool-users in the ontology corpus?’). More complex queries involve reasoning over the structure of the instances themselves, to answer more general questions such as:

> *What are the differences between hominin bone marrow extraction and primate nut extraction?*

The answer requires the comparison of matching structures in the ontology. For example, the concept of ‘extraction’ or of ‘accessing nutritious content from a hard container’ are form matches. The design of efficient ontology-matching algorithms to mine data by comparing bouts in a structural manner as we have described above is a challenge. In Figure 11, we provided a possible pattern that could be searched within the ontology to compare the action that consists in breaking open a container with a stone tool to access its contents. In the case of the observed behaviour of macaques, concepts A and B are then matched with *Nutshell* and *Nut*, respectively. Our comparison could be tool SHK-2152, a stone used in percussive activity at Olduvai Gorge (de la Torre et al. 2013; Arroyo and de la Torre 2016; Leakey 1971; Arroyo and de la Torre 2016). Here concepts A and B in Figure 11 are matched with *Bone* and *BoneMarrow*, respectively.

**Figure 11:**
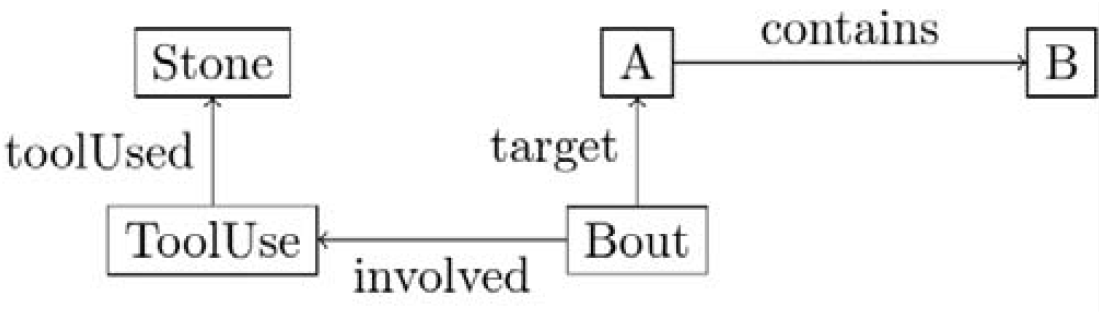
Pattern for breaking open a container with a stone tool.

Instances of tool use that contain this pattern can then be aggregated from the ontology and compared. In Figure 12, this common pattern to match is indicated in red.

**Figure 12:**
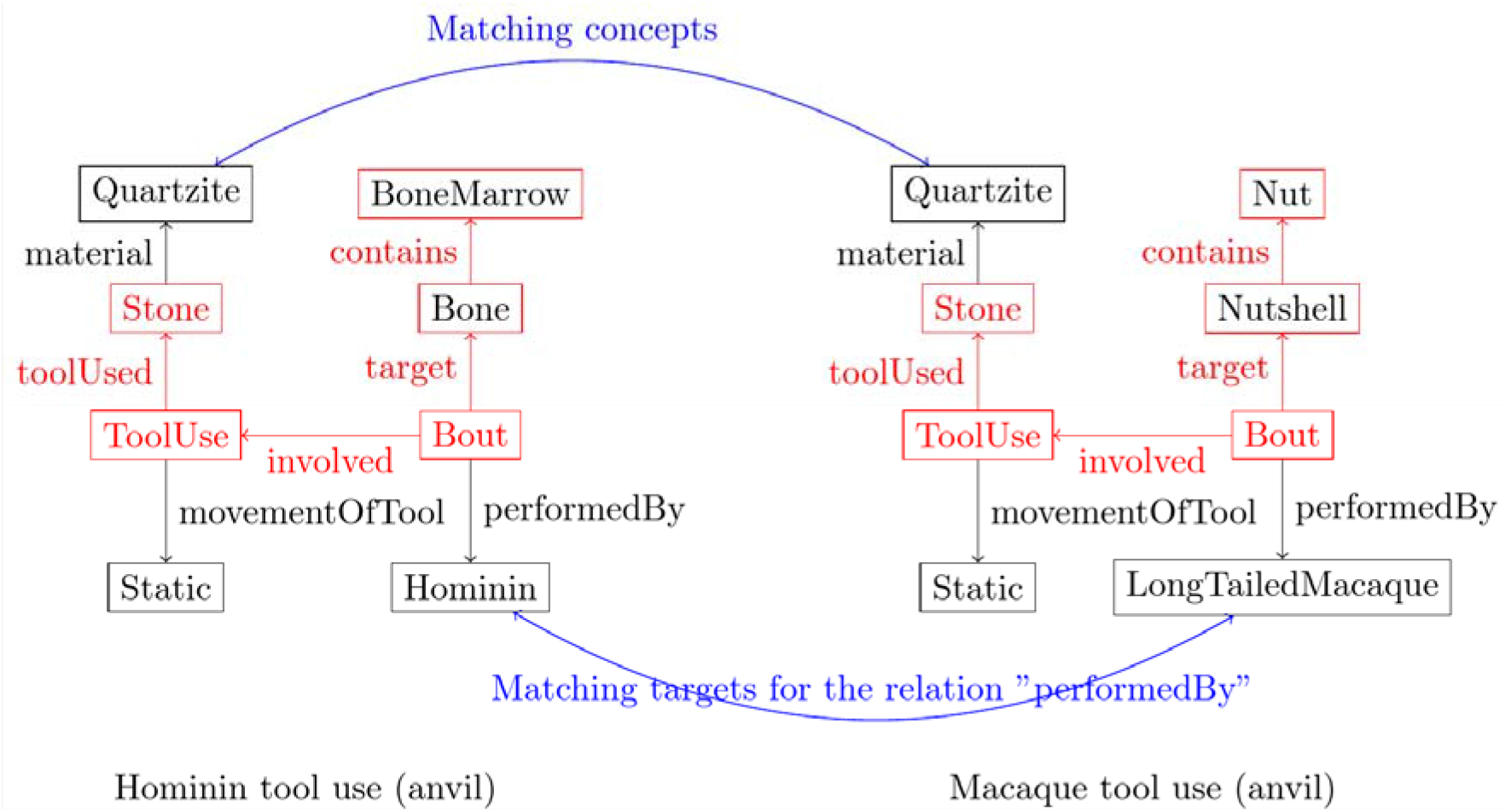
Comparison of two different instances of tool use.

In this example, both tools are made from quartzite, as highlighted with a curved blue arrow labelled ‘Matching concepts’ in Figure 12. These matching concepts indicate that the graph should be made more precise in this area. For example, we might add use-wear analysis from (Arroyo and de la Torre 2016). The problem with these matching concepts is that they are highly dependent first on the quality of the reconstitution and the descriptions, and second on having a harmonized structure which we discuss below. The assumption that we can match concepts also need to be explicit and verified. Another match is in the target of the relation *performedBy*: who is using the tool? Finer description is not necessary here, because details such as motion or anatomical characteristics can already be captured by the rest of the graph. With complex structure analysis we can identify matching behaviours based on their structure. This is an additional feature that *chaînes opératoires* cannot offer and it is a great aid when making large-scale comparisons or sub-datasets using the data in the ontology.

#### Inferring actions from instances

This challenge is related to both the problem of reconstruction in archaeology and the problem of *decomposition* of *behaviour into actions*. Generally, we miss information about the actions needed to reach a goal – tool use or tool making. In our example of BT, the honey-dipping chimpanzee, the textual description mentions the actions of inserting a stick into a hole in a trunk, thrusting the stick in a certain manner, withdrawing it, and eating honey that adhered to the stick. In the same text BT is observed to pick-up and modify another stick. In archaeology we do not have access to direct observations of behaviour and entire technological processes must be reconstructed to reach an idealized description of an ancient process, with varying degrees of confidence. It is likely that parts of actions are missing, and critical steps might be missing, inferred wrongly or the complexity of an overall behaviour could be oversimplified. For example, the difficulty of procurement of a tool may have repercussions on the choice of tool in the overall behaviour, but if the procurement step is not indicated in a material study, this difficulty will not appear, or it is difficult to deduce. One possible solution is to record the nature of the behaviour in the ontology, whether it is reconstituted and by whom, directly observed, or inferred by a machine learning algorithm, with an explicit measure of the degree of confidence.

#### Handling incomplete data

The problem of incomplete data is related to the previous question of inferring actions with a certain degree of confidence, especially in extant archaeological behaviours. However, primatology and observations of behaviour also may lack information, and from a conceptual standpoint there are several solutions, which need to be considered. Modelling of (un)certainty has been explored in AI and robotics and may provide a solution here (Russel and Norvig 2020, chapter 13). For example, concepts that can express the lack of knowledge can be added: *UndeterminateAge, UndeterminateSex*. However, the problem of when to stop describing what we do not know also arises, and a convoluted ontology with ‘empty’ concepts becomes unreadable. Another solution is to infer knowledge much like we might do for actions, and ‘fill up the holes’ in the descriptions. However, there needs to be a measure of the quality of the inferences, as we mentioned above.

#### Finding raw data for tool use descriptions

The data in the ontology comes from our tailor-made corpus, so any bias from the corpus also appears in the ontology. For example, the corpus contains articles that consist of meta-information, conclusions, and interpretations of tool uses. These sources provide information on how many tools were successfully used by a group of primates over a period, and important vocabulary and taxonomical relationships. However, they contain no qualitative description of these successful tool-use bouts. If we use statistical interpretations, we risk missing, for example, outliers that prove capacities for certain types of tool handling and relevant details on tool use and making are lost. This “lack of statistical significance” explains how, for example, many deception tactics in primates – wherein a primate will try to mislead another on purpose – were ignored and went unpublished before Byrne’s survey work (Byrne 1995).

It is therefore vital to mine the rawest data available. Those are descriptions of specific instances of tool use, for example the descriptions of tool use events captured via camera traps in the supplementary material in original sources (like Bessa, Hockings, and Biro 2021). This would be reproduced in an extract as follows: “Adult female is seen dangling from tree branches (…), breaks twig with right hand while holding herself with left, inserts proximal end into hive entrance, brings it to mouth, discards tool, rapidly descends and runs away”. Such a textual description is immediately useable and easy to describe in the ontology. In its current state, the ROCA ontology contains only data obtained from such textual descriptions.

#### Redundancy of information and multiple ways of saying the same thing

In some cases, the same information can be stored in different parts in the ontology. For example, the concept of *EdgeHammering already* includes information on the way the tool is being handled. In addition, it could also appear as a property of the hammer itself during the bout as it gives information on its angle during hammering. For our purpose it is not a problem if the information appears more than once, because the focus was on the recording of information. However, a normalized and unified way of describing tool use bouts that verifies this criterion of non-redundancy is needed, much like has been done in formal logic (Crama and Hammer 2011). Non-redundancy of information in a representation has positive effects on both human readability and computational complexity. For example, while on one hand redundancy increases reliability in computers systems, it also requires more memory power. Related to this problem is the idea that the same event can be described in different ways. For example, the repetition of the movement of hammer-use may be described by listing the repeated events (hammer use, then hammer use, repeated a certain number of times) or by indicating explicitly how many times the event was repeated. Different concepts may also be captured in different manners. One solution is to subdivide as much as possible. For example, consider one concept for a chimpanzee from *Mahale, MahaleChimpanzee*, and two different concepts *Chimpanzee* and *Mahale*. Because there could be a relation of provenance, we choose the latter notation for reusability and modularity’s sake and now the same concept can be used in different contexts.

### 4.3. Chaînes opératoires and ontologies

In the introduction, we identified several differences between *CO* and an ontology approach. First, the ontology makes no assumption towards a known outcome, unlike *CO*. Second, concepts, connections, and assumptions are made explicit in an ontology, which aids comparability and knowledge discovery. Third, the language we use is grammatically strict. There are specific syntactic guidelines, defined by the conventions for OWL, the language we use to express the concepts, the relations, and the instances in our ontology. The concepts used across all our instances of behaviour are also the same. This means that representations within the same ontology are unified and can be compared with each other automatically.

Ontological representations are also at least as powerful, in terms of representation, as *CO*. In other words, whatever can be described with *CO* can be described with an ontology. In the context of lithic analysis, (Bar-Yosef and Van Peer 2009) describe the nature of the *CO* approach. Syntactically, that is, when considering what is needed to express *CO*, the approach is characterized by several features that can be theoretically captured by ontologies. First, any concept in natural language can be defined within the ontology. This makes it possible to describe gestures, central to tool or technology analysis (Schlanger 1990, p. 20), objects, and abstract concepts used in *CO*. Second, the problems of representing “sequence of technical actions” (Tixier, Inizan, Roche, 1980, p. 8), “chronological, sequential representation” (Geneste, 1989, p. 76-77), or “placing each piece [of an assemblage] in a reduction sequence” (Delagnes 1995, p. 202, citing Pelegrin 1986) are answered using temporal relations between actions, or between in the ontology. Third, a “classification system” approach (Geneste, 1989, p. 76-77) is answered by using taxonomical relations in the ontology.

We have verified experimentally that *CO* can be represented without loss of information using an ontology by decomposing a *CO* for pottery production from (Gosselain 2018) (Figure 13). We gave the *CO* an ontology-based representation with a taxonomy of the relevant concepts.

**Figure 13:**
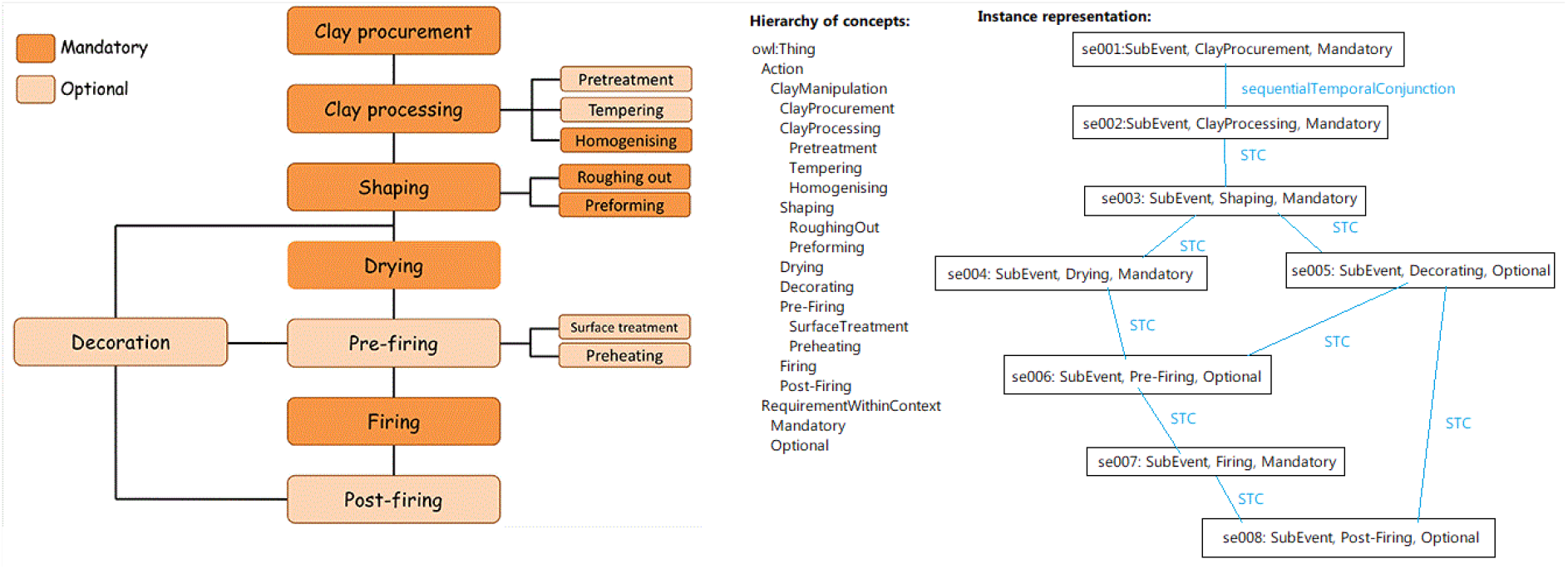
Pottery *chaîne opératoire* (reproduced from Gosselain 2018) and a suggested ontology-based representation.

## Conclusion

We set out to contribute to current discussions on the complexity of extinct and modern non-human and human primate tool tasks. We thus explore ontologies as a useful means of representing tool use and tool making that can help with perceived shortcomings of common representation schemes such as textual descriptions. Ontologies, that seen as taxonomies with more descriptive relations between concepts, are widely used in linguistics, medicine, archaeology, and cultural heritage as a mathematical basis for databases. This helps experts organize, reason on, and discover new knowledge related to their fields. In archaeological, anthropological, and primatology literature, observations or reconstructions of tool use are typically organized in chaînes opératoires or textual narratives. This hampers both comparisons and automated knowledge discovery because there is no common scheme of description, and it is difficult to aggregate data described in schemes that vary from one researcher to another. In particular, the comparison of behaviours remains superficial.

We set out to create a unifying ontology for primate tool use, named ROCA. First, we built a representative corpus of articles and books on primate tool use and tool making (sources N = 75). Then, we extracted and provided semantic structure such as taxonomical relationships to relevant vocabulary, both manually and automatically, for the latter using state-of-the-art NLP text mining techniques. For example, we give physical concepts to describe nut-cracking behaviour alongside a taxonomy of nuts. The ROCA ontology (https://rocaontology.github.io/) contains 900 concepts to describe physical movements down to the anatomical level, tools and their materials, ecology, and primate features. It also contains concepts and relations that allow us to describe behaviours in a graph-like manner, adding to the ease of interpretation. For example, we define sequential relations to tie up “picking up a rock, placing a nut on an anvil, hitting the nut, eating the nut” in the order in which it has been observed.

Due to its nature as an ontology and our experiments, we argue that ROCA has the following qualities. First, the data is uniform because the instances of behaviour are described using the concept and the guidelines provided by the ontology. Every instance is expressed using the same formal language (OWL). Second, the data is unified: no matter the source, everything is stored within the same database. This makes querying and knowledge discovery easier. Third, the data can be modified or added upon easily depending on the needs of the researcher. We have been continuously adding new data in the ontology. Fourth, the ontology is acentric: the researcher can choose their focus and filter results according to their needs. Where we would need at least two taxonomies for a study of chimpanzees involved in nut cracking - that is, a taxonomy for chimpanzees and a taxonomy for nuts-the ontology contains both and the centre can be chosen. Fifth, ROCA is human-readable. The concepts are expressed in natural language, can be provided a definition, and are given a graphical representation to aid at interpreting the behaviour.

We prove the viability of the ontology in helping knowledge discovery. We perform several queries and prove that we can perform meta-analysis on the data by answering questions such as “what different types of hammering are there?”, “How many cases of female tool use are there in the ontology?” (126 cases) and “Can we conclude that an individual is male, or female based on their tool use behaviour?” (No).

Areas of improvement include an assessment of the useability of the ontology amongst researchers, of the quality of the instances, of the statistical significance of the instances, and of the consistency of concepts depending on their sources. We conclude with some planned work: creating a matching algorithm to delve deeper into the comparison of instances, shedding light into the decomposition of behaviour into actions, the inference of reconstituted tools from actions, and the handling of incomplete data.

We finally show that chaînes opératoires, which are a typical and common way of representing behaviour, can be represented within an ontology with all the benefits that it entails and that we listed above, thereby approaching CO in a different light. For example, we concentrate on automated comparison for automated knowledge discovery by aggregating the data we obtain with unified syntactic guidelines. We also avoid complex conceptual assumptions such as deciding on a direction towards a known outcome, by focusing solely on the problem of describing and recording actions for the purpose of later analysis.

## Acknowledgements

This work was conducted thanks to the support of the Faculty of Mechanical Engineering of Delft University of Technology through the ROCA Cohesie grant. In addition, this project was also funded by the European Research Council under the European Union’s Horizon 2020 research and innovation program under grant agreement 804151 (grant holder G. L.).

The authors would like to express their gratitude to Dr. Sebastian Fajardo for his insightful comments on earlier versions of the manuscript.

## Footnotes

1 The domain of the relation is the set of values it can take as input; the codomain is the set of outputs. In the example that follows, *describedInText* has for domain a set of behavioural instances and for codomain a set of texts.

## Notes

https://rocaontology.github.io/

### Competing Interest Statement

The authors have declared no competing interest.

### Summary of Updates

Misc. information regarding chaines operatoires. Better coherence within abstract and conclusion.

